# Distinct Amygdala–Frontopolar and Amygdala–Medial Temporal Pathways Support Temporal Memory for Emotional Events

**DOI:** 10.64898/2026.02.19.706842

**Authors:** Jingyi Wang, Joanne E. Stasiak, Neil M. Dundon, Elizabeth J. Rizor, Parker L. Barandon, Christina M. Villanueva, Scott T. Grafton, Regina C. Lapate

**Author notes:** Corresponding authors: Regina C. Lapate or Jingyi Wang. **Conflict of Interest Statement:** The authors declare no competing financial interests.

## Abstract

Growing evidence suggests that emotion shapes temporal aspects of memory—such as remembering when an event occurred—yet the neural bases of these effects remain unclear. Prior work indicates that temporal-context representations in memory are supported by the function of medial temporal lobe (MTL) and prefrontal regions, including hippocampus, perirhinal cortex, and lateral frontopolar cortex (FPl). These regions have privileged access to amygdala-originated affective signals that may mediate emotional influences on temporal memory. To test whether these candidate pathways shape temporal memory for emotional events, participants encoded trial-unique negative and positive images in the scanner, followed by a surprise temporal memory task. We found that more negative stimuli were associated with greater amygdala activation and lower temporal-memory error for when emotional images occurred. Replicating and extending prior work, higher stimulus-evoked multivoxel pattern similarity in the hippocampus and FPl was associated with higher temporal memory error. Notably, greater amygdala responding to emotional images predicted higher between-trial pattern similarity in the hippocampus, but lower within- and between-trial similarity in FPl. Amygdala engagement was also correlated with greater perirhinal activation, which in turn predicted stronger item memory and a recency bias in temporal memory estimates. Collectively, these findings reveal distinct—and opposing—modulatory effects of amygdala–FPl and amygdala–MTL pathways on temporal memory, with the potential to sharpen and blur memory for when emotional events occurred.

## Introduction

Emotional events powerfully influence how we remember time (Clewett & Murty, 2019; Palombo & Cocquyt, 2020; Talmi & Palombo, 2025; Wang et al., 2022), an essential feature of episodic memory. For example, one may vividly recall the precise sequence of a door opening, friends shouting, and the reveal of a cake at a surprise birthday party, whereas ordinary moments from earlier that day blur together. A recent surge of behavioral studies have underscored potent yet heterogeneous effects of emotion on temporal memory, with evidence for both enhancement (Bogdan et al., 2023; Cliver et al., 2024; Lake et al., 2016; Palombo et al., 2021; Riegel et al., 2023) and distortion (Colson et al., 2025; Laing & Dunsmoor, 2025; Laing et al., 2026; McClay et al., 2025; Wang & Lapate, 2025) of temporal memory by emotion. Despite the pervasive and occasionally divergent influence of emotion on temporal memory, the neural mechanisms underlying their interplay remain poorly understood.

Existing evidence suggests that function of the hippocampus, lateral frontal pole (FPl), and the entorhinal cortex supports temporal memory (reviewed in (Buzsáki & Tingley, 2018; Clewett et al., 2019; Tsao et al., 2022)). For instance, greater similarity of neural activity patterns in response to distinct events in the hippocampus has been linked to shorter objective (Hsieh et al., 2014; Nielson et al., 2015; Wang et al., 2025) as well as remembered (Ezzyat & Davachi, 2014) temporal distances. Moreover, higher between-trial pattern similarity in the hippocampus (Jenkins & Ranganath, 2016; Manns et al., 2007) and FPl (Jenkins & Ranganath, 2010) has also been linked with poorer temporal memory fidelity (e.g., reduced temporal order memory accuracy) (but see (DuBrow & Davachi, 2014)). However, these studies have primarily examined how relatively neutral events (e.g., objects) are organized in memory, leaving it unclear whether and how these candidate neural circuits similarly support temporal memory for emotionally evocative events, as well as whether they interact with emotion-coding regions during emotional processing.

The amygdala—a set of subcortical nuclei subserving the early appraisal of emotionally relevant stimuli and the orchestration of emotional responses (LeDoux, 2007; Paz & Pare, 2013; Tye, 2018)—has strong direct anatomical projections to the hippocampus (Wang & Barbas, 2018) and perirhinal cortex (Stefanacci et al., 1996), and multisynaptic projections to FPl (Ghashghaei et al., 2007; Medalla & Barbas, 2010). The amygdala and perirhinal cortex closely interact during emotional-event processing, with perirhinal engagement typically associated with enhanced item memory (Bisby & Burgess, 2017; Ritchey et al., 2019). In a separate line of work, perirhinal engagement putatively supporting item strength has been proposed to produce recency biases in temporal memory judgments (DuBrow & Davachi, 2017; Jenkins & Ranganath, 2016).

Recent work indicates that strong amygdala-hippocampal projections can exert a powerful and long-lasting influence on hippocampal neurons, including the potential suppression of entorhinal-hippocampal inputs thought to be essential for intact hippocampal temporal coding (Eichenbaum, 2014; Robinson et al., 2017), via inhibitory parvalbumin neurons (Wang & Barbas, 2018). This, in turn, can result in less discriminable hippocampal patterns over time, and lower-fidelity coding of temporal context (reviewed in (Wang et al., 2022)). In contrast, the connectivity profile between the amygdala and FPl suggests that amygdala-frontopolar interactions—likely mediated by a polysynaptic pathway via ventromedial prefrontal cortex (vmPFC; BA 25 and BA 32) targeting upper layers of FPl and innervating inhibitory calbindin neurons (Joyce & Barbas, 2018; Medalla & Barbas, 2010)—have the potential to sharpen FPl-supported representations during emotional processing (Barbas et al., 2018).

In the current study, we tested whether these previously identified neural correlates of temporal memory—including hippocampal and FPl pattern similarity and perirhinal activation at encoding—support temporal memory for emotional events, and whether they are modulated by amygdala responses to emotional pictures. Participants encoded trial-unique positive and negative images in the MRI scanner. After finishing the scan, participants completed a surprise memory test, where they indicated whether they remembered each image as well as *when* during the MRI experiment they thought each emotional image had been shown (**Figure 1A-C)**. First, we tested whether picture valence and arousal influenced temporal memory accuracy using normative ratings. Next, we examined whether temporal memory accuracy varied as a function of pattern similarity in the hippocampus and FPl, estimated both between trials (Jenkins & Ranganath, 2010) as well as within trials (Sinclair et al., 2021). Then, we tested whether amygdala responses to emotional images modulated these similarity measures and the magnitude of perirhinal engagement. Given previous evidence showing that higher hippocampal pattern similarity is correlated with lower temporal memory accuracy (Jenkins & Ranganath, 2016; Manns et al., 2007), we hypothesized that greater pattern similarity in the hippocampus and FPl would be associated with poorer temporal memory, and that amygdala activation in response to emotional pictures would modulate pattern similarity in these regions. We further hypothesized that perirhinal activation during emotional-picture processing would covary with amygdala activation and be associated with recency biases in temporal memory estimates.

**Figure 1.**
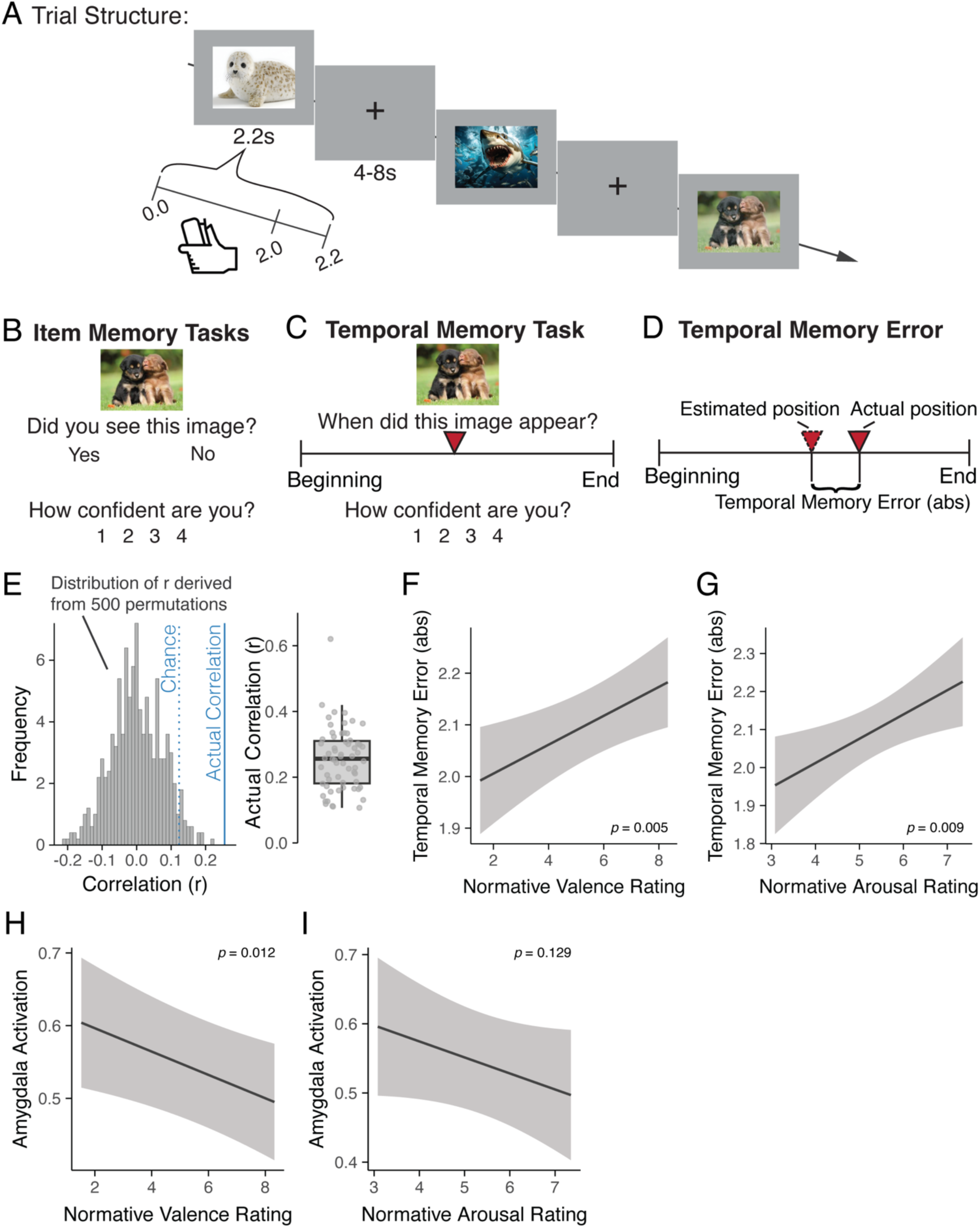
Experimental Design and behavioral results. **A.** The trial structure of the scanner task is shown. Trial-unique negative and positive images were presented for 2.2 s. During image presentation, participants made a joystick response according to a rule provided at the beginning of each block (see *Methods*). **B-C.** After completion of the scanner task, participants completed a surprise memory task. Each trial began with an item memory judgment in which participants indicated whether or not they remembered seeing each item during the scanner task (‘Yes’ or ‘No’), followed by a confidence rating ranging from 1–4 (‘1’: ‘not at all confidence’, ‘4’: ‘very high confidence’) (B). Participants were then asked to indicate, using a continuous timeline slider, *when* they thought the image had been presented during the scanner task, followed by a confidence rating (C). **D.** Temporal memory errors were computed as the absolute difference score between each image’s remembered and actual presentation times during the session. **E.** Non-parametric permutation test of temporal memory showing an above chance correlation between estimated and actual timeline positions for an example participant (left). Dotted blue line: chance level as 95% percentile, blue line: correlation. Boxplot: the distribution of correlations between actual and remembered picture presentation times across participants (right). **F-G.** Temporal memory errors were positively associated with emotional valence (F), and positively associated with emotional arousal (G). **H-I.** Amygdala activation was negatively associated with emotional valence (H), but not reliably associated with emotional arousal (I). Shaded ribbon in F-I: 95% confidence interval (CI).

## Materials and Methods

### Participants

Seventy-seven participants were recruited from Santa Barbara, CA (mean age = 20.579 years; SD = 1.627; range = 18 – 26 years; 56 female; 41.0% White, 31.3% Asian, 22.9% Hispanic/Latino, 1.2% Black, and 3.6% other races). Six subjects were excluded due to excessive motion (> 1 mm) or difficulty understanding task instructions, which yielded a final sample of seventy-one participants (mean age = 20.557 years; SD = 1.557; range = 18 – 26 years; 51 female; 42.5% White, 30.1% Asian, 24.7% Hispanic/Latino, 1.4% Black, and 1.4% other races). All participants were healthy, with normal or corrected-to-normal vision, and no self-reported history of psychiatric or neurological disorders. Written informed consent was obtained from every participant. Participants were recruited at the University of California (UC), Santa Barbara. All study procedures were approved by the UC Santa Barbara Human Subjects Committee. Participants were compensated monetarily for their participation.

### Statistical power

We based our power analysis on prior findings showing an association between between-trial pattern correlation in the FPl and hippocampus and temporal memory accuracy. From these findings, we derived conservative effect sizes of f = 0.32 for FPl (Jenkins & Ranganath, 2010) and f = 0.57 for hippocampus (Jenkins & Ranganath, 2016). An a priori power analysis conducted in G*Power 3.1 (Faul et al., 2007) indicated that n = 22 participants would provide > 80% power at α = 0.05 to detect effects of these magnitudes in both regions. Given our final sample of N = 71, a post-hoc power analysis indicated that the current study had > 99% power to detect these effects in both regions. Thus, our sample was adequately powered.

### Image stimuli

A total of 192 images were selected from the International Affective Picture System (IAPS, (Lang et al., 2008) and Open Affective Standardized Image Set (OASIS, (Kurdi et al., 2017). We selected 96 negative stimuli (*M*_valence_ = 3.207, SD_valence_ = 0.867; *M*_arousal_ = 6.019, SD_arousal_ = 0.632) and 96 positive stimuli (*M*_valence_ = 7.448, SD_valence_ = 0.502; *M*_arousal_ = 4.698, SD_arousal_ = 0.636) balanced across social and nonsocial categories using the IAPS standardized ratings for emotional valence and arousal (where 1 = very negative/low-arousal, and 9 = very positive/high-arousal). Valence and arousal ratings for OASIS images were converted to IAPS’ standardized ratings (rating_OASIStoIAPS_ = ((rating_OASIS_ -1)*8/6) + 1). Negative images were significantly more negative (*t* = -41.47, *p* < 0.001) and arousing (*t* = 14.44, *p* < 0.001) than positive images. To disentangle the potentially independent contributions of emotional valence and arousal to temporal memory, all analyses in the present study used continuous normative valence and arousal scores as simultaneous regressors in a multiple regression model predicting temporal memory.

### Procedure

#### Overview

Following informed consent and MRI safety screening, participants received instructions for the encoding task. They then completed a high-resolution T1-weighted anatomical scan, followed by approximately 40 minutes of task fMRI data collection with trial-unique emotional image stimuli. Immediately after the completion of the task, participants underwent a resting-state scan (*data not reported here*). Upon scanning completion, participants performed a surprise memory task in which they indicated whether they remembered each previously-presented image (item memory), as well as *when* during the MRI session they thought each image had been shown (temporal memory). Both types of memory judgments were followed by confidence ratings, as detailed below.

#### Experimental design

##### Encoding task

In the MRI scanner, participants completed an Approach-Avoidance task (adapted from (Bramson et al., 2020)), which consisted of eight functional runs (lasting ∼ 4 minutes each). The approach-avoidance manipulation was included as part of a larger study investigating emotion-motor interactions and was not a focus of the present study. We therefore balanced motor goal across emotional valence conditions and controlled for this factor in all of the relevant analysis models, as detailed below.

On each trial, a trial-unique emotional image was presented for 2200 ms. Participants performed a joystick movement within the first 2000 ms of image presentation (*M*: 1309 ms, *SD*: 112 ms) according to a rule (“approach positive, avoid negative” *vs*. “approach negative, avoid positive”). Following participants’ joystick responses, images were enlarged (following approach) or reduced (following avoidance) by 30% of their original size until the end of the trial. Action goal was manipulated orthogonally in relation to the emotional valence manipulation. Each run contained two rule blocks (comprising 12 trials/each): “approach-positive + avoid-negative” and “approach-negative + avoid positive”, the order of which was counterbalanced across runs. Rule instructions were displayed for 7s at the beginning of each block. After each trial, an intertrial interval followed (4, 6, 8 seconds sampled from a quasi-exponential distribution). Image presentation order was pseudorandomized across participants, ensuring that images with the same emotional valence or action goal were not presented consecutively more than three times. As detailed below, all temporal memory analyses controlled for participants’ trial-wise joystick action, and the reported results did not interact with trial-wise action goal.

##### Surprise memory tasks

Upon completion of the MRI scan, participants performed item and temporal memory tasks outside the scanner (time between scan completion and the start of the surprise memory tasks: *M* = 11.89 min, SD = 3.58 min). In each trial, participants viewed an image previously presented during the encoding task, with trial order randomized. Participants first judged item memory by answering “*Did you see this image before?*” with a binary “Yes”/“No” choice (**Figure 1B)**, followed by a confidence rating (“*How confident are you in your response?*”) on a scale of 1 *(“Not at all confident”)* to 4 *(“Very confident”)*. Next, they completed a temporal memory task in which they estimated *when* each image had appeared during the experiment (“*When did you see this image during the task?*”) using a timeline slider spanning the beginning to the end of the task, followed by a confidence rating (**Figure 1C).** Confidence ratings for temporal memory judgments are reported in supplementary materials *(Supplementary Materials: confidence rating analyses,* **Supplementary Figure 5**). Responses were self-paced, with a 500 ms intertrial interval.

### Functional MRI Methods

#### Image acquisition

Neuroimaging data were acquired in the UC Santa Barbara Brain Imaging Center with a Siemens Prisma 3T MRI scanner equipped with a 64-channel radio frequency head coil. Whole-brain Blood Oxygenation Level-Dependent (BOLD) fMRI data were obtained using a T2*-weighted 2-accelerated multiband echoplanar imaging sequence (54 axial slices, 2.5 mm^3^ isotropic voxels; 80 x 80 matrix; TR = 1900 ms; TE = 30 ms; flip angle = 65°; 133 image volumes/run). High-resolution anatomical scans were acquired using a T1-weighted magnetization-prepared rapid gradient echo (MPRAGE) sequence at the beginning of the session for spatial normalization (TR = 2500 ms; TE = 2.2 ms; flip angle = 7°), followed by a gradient echo field map (TR = 758 ms; TE1 = 4.92 ms; TE2 = 7.38 ms; flip angle = 60°).

#### Regions of interest

##### Subcortical

Hippocampal and amygdalar regions of interest (ROI) were defined using the Harvard-Oxford subcortical structural atlas (Desikan et al., 2006; Frazier et al., 2005) thresholded at 50%. Each MNI space mask was registered to participants’ anatomical space using FNIRT (10 mm warp resolution) while maintaining functional resolution (2.5 mm^3^).

##### Cortical

The perirhinal ROI (BA35) was derived from FreeSurfer (Augustinack et al., 2013). The FPl ROI was obtained from the Oxford PFC Consensus Atlas (Sallet et al., 2013) (http://lennartverhagen.com/) in FreeSurfer space. Atlas masks were then thresholded at 25% and registered to participants’ native surface space using FreeSurfer. Then, vertex coordinates for the perirhinal and FPl masks were transformed into each subject’s volumetric space. ROI masks in volumetric space were constructed by projecting half the distance of the cortical thickness at each vertex, with functional voxels required to be filled at least 50%. Volumetric masks were then resampled to functional resolution (2.5 mm^3^).

### Statistical analysis

#### Temporal memory error score and behavioral analyses

Participants used a timeline slider to indicate *when* they thought each image was presented during the experiment. We quantified trial-wise temporal memory errors by taking the absolute difference between each image’s estimated and actual positions on the timeline slider (**Figure 1D**; (Montchal et al., 2019)). Image presentation times (ranging from 27.512s to 3706.315s from the beginning to the end of the scanner task) were converted to a run-time scale (0.000–8.000) using the following formula: image presentation time * (8 / total session time), thereby normalizing participants’ individual timelines and maintaining three-decimal resolution.

To assess temporal memory performance at the subject level, we generated a null distribution by randomly shuffling temporal memory estimates across trials for each participant (n=500 times) and correlating each permuted set with the true temporal positions (for an example, see **Figure 1E**). Participants’ observed correlation between estimated and true temporal positions was compared with their own permutation-derived null distribution to derive statistical significance. Sixty-two (out of 71) participants performed significantly above chance (mean age = 20.63 years, SD = 1.60, range = 18–26 years; 45 female, **Figure 1E**) and were retained for temporal memory analyses. For the analyzed sample, the average correlation between actual and estimated timeline position was *r* = 0.256 (*SD* = 0.09).

To examine whether and how emotion influenced temporal memory, we fitted normative image valence and arousal ratings to predict temporal memory error using mixed-effects models in R (lme4 package; (Bates et al., 2015)). We controlled for action goal (i.e., instructed joystick movement direction: approach (forward) vs. avoid (backward)) in these mixed-effects models. Participants were modeled with random intercepts, and valence ratings were modeled as random slopes.

#### fMRI data modeling

##### fMRI data preprocessing

Functional neuroimaging data were preprocessed using FEAT (FMRI Expert Analysis Tool) version 6.00, implemented in FMRIB’s Software Library (Jenkinson et al., 2012; Smith et al., 2004). Preprocessing steps included applying high-pass temporal filtering with a 100s cutoff, FILM correction for autocorrelation in the BOLD signal, motion correction using MCFLIRT, slice-time correction, and creation of a confound matrix with points of framewise displacement changes larger than 0.5 mm to be used as regressors of non-interest to control for movement-confounded activation. Data were smoothed using a 3 mm full-width at half-maximum Gaussian spatial filter. Brain extraction was carried out through Advanced Normalization Tools (ANTs) skull-stripping algorithm (Avants et al., 2009). Functional images were co-registered to participants’ T1-weighted anatomical image using a linear rigid body (6-DOF) transform while maintaining native functional resolution (2.5 mm^3^ isotropic).

##### Single-trial GLM

We obtained voxel-wise and trial-wise BOLD activation parameter estimates using the Least-Squares All general linear model (GLM) approach (Mumford et al., 2012) in FEAT models in FSL (Jenkinson et al., 2012). Single trials were modeled using a canonical hemodynamic response function (Double γ). Then, we applied multivariate noise normalization to the parameter estimates (Walther et al., 2016). Specifically, we estimated the noise covariance from the residuals of the GLMs for each ROI, regularized this matrix using the optimal shrinkage parameter, inverted it, and multiplied it by the vector of trial-wise betas (Ballard et al., 2019; Lapate et al., 2022). This procedure has been shown to reduce nuisance correlations between voxels caused by physiological and instrumental noise (Walther et al., 2016).

##### Single-trial univariate analysis

To estimate univariate activation at the trial level, voxel-wise parameter estimates were averaged within each ROI (here, amygdala and perirhinal cortex).

##### Between-trial pattern correlation

Pattern correlation between trials was calculated as the Pearson correlation of trial-wise neural activation patterns (i.e., parameter estimates) between each trial and its temporally-adjacent neighbors (trial lags ± 1, 2, 3, 4) for each ROI (here, hippocampus and FPl). Pearson correlations were Fisher’s Z transformed.

##### Within-trial pattern correlation

Given recent results suggesting that contextual changes known to sculpt temporal memory (i.e., event boundaries) alter pattern correlation within trials (TR-by-TR; also termed temporal autocorrelation) in regions involved in temporal memory such as the hippocampus (Sinclair et al., 2021), we computed the within-trial pattern correlation from the preprocessed functional data (see Methods: *fMRI data preprocessing)*. To do so, we extracted voxel-wise activity at each TR within each *a priori* ROI, which were then z-scored per voxel, run, and participant. Next, we correlated the z-scored activation patterns at timepoints *T* and *T+1* (per ROI). These within-trial pattern correlation values were then standardized (Fisher’s Z transformed). To measure the persistence of neural activation patterns into the stimulus offset period (Puccetti et al., 2021), we focused our primary analyses on the activation pattern correlation between TRs 3 (corresponding to BOLD responses during picture presentation) and 4 (corresponding to BOLD responses after picture offset) after picture onset (corresponding to 5.7 – 7.6s from stimulus onset, **Figure 2B**). To ensure that observed effects putatively indexing representational persistence were not merely due to the intrinsic autocorrelation of the BOLD signal, we re-ran these analyses using the residual time series obtained from the single-trial GLM.

**Figure 2:**
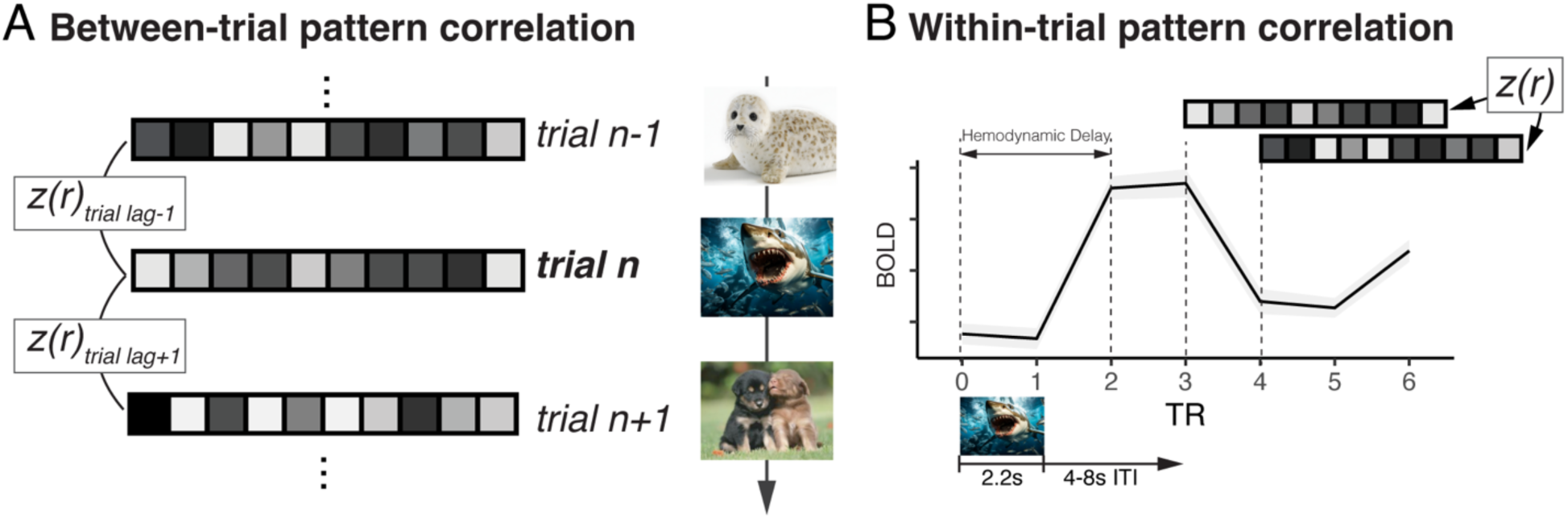
Multivariate pattern analysis approach. **A.** Between-trial pattern correlations were obtained for each ROI by computing the correlation of multivoxel activity patterns (derived from parameter estimates from single-trial GLMs, see *Methods* for details) between a given trial and its adjacent trials. **B.** BOLD activity patterns were extracted from each ROI at each TR following image onset and correlated between TR pairs to obtain within-trial pattern correlations. Bottom image: image onset and offset are shown in relation to the TRs examined. ITI: inter-trial-interval.

#### Neural-behavior correlations

##### Between-trial neural pattern correlation analyses

We examined the relationship between between-trial neural pattern correlation and temporal memory using categorical and continuous estimates of temporal memory in mixed-effects models. First, to more directly reflect the approach used in previous studies, we first examined temporal memory accuracy using a categorical approach (Jenkins & Ranganath, 2010; Montchal et al., 2019; Zou et al., 2023), whereby temporal memory performance scores (per participant) was divided into tertiles: ‘High, ‘Medium’, and ‘Low’. Then, we tested whether temporal memory performance (High *vs.* Low) was associated with the magnitude of neural pattern correlation between each trial and its adjacent neighbors (trial lags ± 1, 2, 3, 4) (Jenkins & Ranganath, 2010). For analyses using continuous estimates, mixed-effects models tested whether between-trial neural pattern correlation at trial lag 1 (i.e., the correlation between patterns in a given trial (*n*) and the following trial (*n+1*)) predicted a given trial’s (*n*) temporal memory error.

We controlled for the following factors in these mixed effects models: normative valence ratings, normative arousal ratings, action goal, whether or not there was a change in valence (same vs. different) or action goals (same vs. different) between consecutive trials, as well as semantic similarity between consecutive image pairs. Of note, semantic similarity has been shown to influence temporal distance estimates as well as neural pattern similarity (Riberto et al., 2022, 2019; Wang & Lapate, 2025). Therefore, we derived semantic similarity estimates between image pairs using deep convolutional artificial neural networks (ANNs) as implemented previously (Kubilius et al., 2019; Simonyan & Zisserman, 2014; Wang & Lapate, 2025). Participants were entered as random intercepts, and normative valence and action goal were entered as random slopes in these mixed-effects models.

##### Within-trial pattern correlation analyses

To test whether within-trial pattern correlation was associated with temporal memory, we performed mixed-effects models with categorical and continuous temporal estimates. First, we tested whether within-trial pattern correlation estimated between each TR pair after stimulus onset (i.e., TR0-TR1, TR1-TR2, TR2-TR3, TR3-TR4) predicted categorical temporal memory performance (High *vs.* Low, as described above), FDR-corrected for multiple comparisons. We also tested whether within-trial pattern correlation at TR3-TR4 in a given trial (*n*) predicted the temporal memory error for that trial (*n*). In these models, we controlled for action goal, valence, and arousal ratings. As before, participants were modeled with random intercepts, and valence and action goal were modeled with random slopes.

##### Perirhinal univariate analysis

To test whether the magnitude of perirhinal activation predicted temporal memory error as a function of session time, we divided the time course of the session (“session time”) into tertiles: Early, Middle, and Late. Mixed-effects models were performed testing the interaction between trial-wise perirhinal activation and session time to predict temporal memory error, while controlling for action goal, valence, and arousal ratings. Participants were modeled with random intercepts, and valence ratings and action goal were modeled with random slopes.

##### Amygdala univariate analysis

To verify whether and how amygdala responses were modulated by emotion—here, emotional valence and/or arousal—we fit separate mixed-effects models using normative valence and arousal ratings to predict trial-wise amygdala activation. Next, to examine whether and how putative neural correlates of accurate temporal memory were modulated by amygdala activation, we fit separate mixed-effects models predicting between-trial pattern correlation (trial lag 1: trial *n*, *n*+1), within-trial pattern correlation (trial n’s TR3-TR4), and perirhinal activation (trial *n*) from amygdala activation at trial *n*. Model structure, including all control variables, was kept consistent with the corresponding models described above.

##### Item memory analysis

We used a mixed-effects model to examine the association between item memory and perirhinal cortex activation. Emotional valence, arousal and action goal were entered as control factors, participants were modeled with random intercepts, and valence ratings and action goal were entered as random slopes predicting item memory and item memory confidence. To control for the potential influence of item memory on temporal memory (Supplementary Results: *Control analyses: item memory*), we entered item memory ratings as a covariate in the above-described mixed-effects models examining the relationship between temporal memory and neural metrics (including between-trial and within-trial pattern correlation as well as single-trial activation models).

##### Mediation analysis

We examined whether the effects of emotion on temporal memory were mediated via amygdala–MTL and amygdala–FPl pathways. To do so, we conducted mediation analyses through Bayesian modeling with the brms package in R (version 2.22.0; (Bürkner, 2017)), which supports mediation analyses on mixed-effects models with random effects. Prior to running these mediational models, we z-scored (across participants) temporal-memory errors. We used weakly informative default priors centered at 0 with a 2.5 SD (Atlas et al., 2022; Callaghan et al., 2021; Vermeylen et al., 2025). Each model was estimated using four Markov chains of 4,000 iterations (including 2,000 warm-up iterations), with a maximum tree depth of 15 and an adapt-delta of 0.95 to ensure adequate convergence. We report the mean of the posterior distribution as the estimate for the standardized ß weight, together with its standard error (SE), and 95% Bayesian credibility interval (95% CI). In addition, we report the probability of direction (*pd*), which is considered an index of effect existence and reflects the certainty with which an effect goes in a particular direction.

## Results

### Negative emotion sharpens memory for when events occurred

We first tested whether emotional valence and arousal modulated memory for when emotional pictures were presented during the experiment. We found that more negative emotional valence was associated with smaller absolute temporal memory errors—in other words, temporal memory errors were lower for more unpleasant pictures (B = 0.028 (SE = 0.01), F = 7.74, *p* = 0.005, **Figure 1F**). Moreover, temporal memory errors were positively correlated with higher arousal ratings (B = 0.064 (SE = 0.024), F = 6.84, *p* = 0.009, **Figure 1G**). Following previous work (McClay et al., 2023), we also examined whether valence and arousal levels interacted in predicting temporal memory. This analysis revealed a trend-level interaction (B = -0.016 (SE = 0.009), F = 2.74, *p* = 0.098), such that for negatively valenced events, higher arousal was associated with greater errors in estimating when the emotional picture was shown (**Supplementary Figure 1**), consistent with prior findings (McClay et al., 2023). These results indicate that more negative event valence sharpened temporal memory for when those events occurred.

### Between-trial pattern correlation: Higher hippocampal similarity is associated with poorer temporal memory

Prior work indicates that higher similarity between adjacent trials in hippocampus and FPl neural activity patterns is associated with poorer temporal memory accuracy for when an image occurred during the experiment (Jenkins & Ranganath, 2010). Thus, we first examined whether we observed these effects in our experiment. To this end, we ran a mixed-effects model predicting neural pattern correlation from temporal memory performance (i.e., high/low) and trial lag.

Critically, the similarity of hippocampal activity patterns between trials significantly predicted temporal memory accuracy (B = 0.013 (SE = 0.003), F = 21.70, *p* < 0.001, **Figure 3A**), such that higher between-trial pattern correlation was associated with larger temporal memory errors. A numerically similar (but marginally significant) effect was also seen in FPl (B = 0.006 (SE = 0.004), F = 3.26, *p* = 0.071, **Figure 3B**), in a direction consistent with prior findings (Jenkins & Ranganath, 2010). Moreover, the main effect of trial lag on pattern similarity was significant in both the hippocampus (B = -0.173 (SE = 0.007), F = 383.16, *p* < 0.001) and FPl (B = -0.177 (SE = 0.010), F = 310.87, *p* < 0.001)—indicating that the between-trial pattern correlation progressively decreased as the trial lag between trials increased in both regions (**Figures 3A & 3B**). Collectively, these results indicate that higher across-trial similarity in image-evoked neural activity patterns at encoding in the hippocampus is associated with poorer temporal memory.

**Figure 3:**
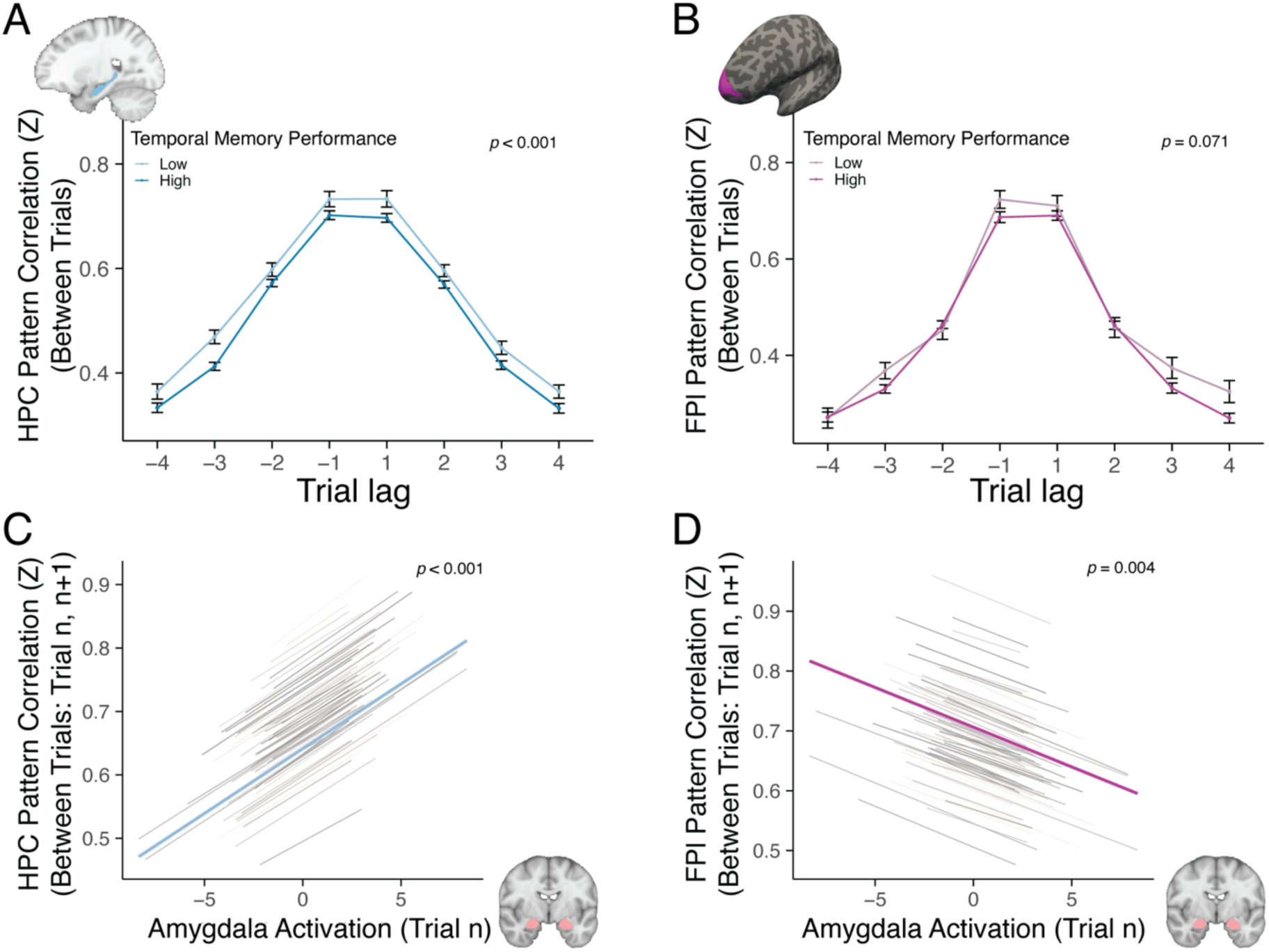
Between-trial pattern correlations. **A** Trials with higher between-trial pattern similarity in the hippocampus were associated with poorer temporal memory performance. **B** A numerical trend was seen in FPl. **C-D** Amygdalar activation was associated with higher pattern similarity across trials (trial lag 1) in the hippocampus (C), but lower between-trial pattern similarity in the FPl (D).

### Amygdala engagement to emotional images differentially modulates hippocampal and FPl between-trial correlation

Higher amygdala activation is often found in response to negative emotional events (Bowen et al., 2018; Kensinger & Schacter, 2006). To determine whether emotional images in the present study reliably modulated amygdala activation, we examined the association between amygdala activation and image normative valence and arousal ratings using mixed-effects models. We found that the magnitude of amygdala activation (across trials and participants) was negatively associated with normative emotional valence ratings (B = -0.016 (SE = 0.006), F = 6.33, *p* = 0.012, **Figure 1H**), indicating stronger amygdalar responses to more negative images, consistent with prior findings (Bowen et al., 2018; Kensinger & Schacter, 2006). Of note, amygdala activation was not reliably correlated with emotional arousal ratings in this sample (B = -0.023 (SE = 0.015), F = 2.21, *p* = 0.129, **Figure 1I**). These results suggest the amygdala activation in this sample tracked emotional valence more reliably than arousal levels. Thus, the analyses that followed focused on the modulation of temporal memory by emotional valence while controlling for emotional arousal (see Methods for details).

Next, we examined whether and how amygdalar engagement during emotional processing shaped the similarity of hippocampal and FPl neural activity patterns over time. We predicted that amygdala activation would exert a prospective effect on hippocampal and FPl similarity, potentially increasing it and ‘blurring’ representations previously associated with temporal memory fidelity (Wang et al., 2022). Consistent with our hypothesis, we found a positive association between amygdala activation at a given trial and subsequent between-trial pattern correlation in the hippocampus (trial lag 1; B = 0.020 (SE = 0.004), F = 31.15, *p* < 0.001) (**Figure 3C**). In contrast, amygdala activation was *negatively* associated with between-trial pattern correlation in the FPl (trial lag 1; B = -0.013 (SE = 0.005), F = 8.45, *p* = 0.004) (**Figure 3D**). Importantly, no significant associations were found between amygdala activation on the current trial and pattern correlation between the current and previous trial in the hippocampus (B = 0.002 (SE = 0.004), F = 0.17, *p* = 0.679) or the FPl (B = -0.003 (SE = 0.005), F = 0.33, *p* = 0.568). Collectively, these results indicate that amygdala activation to emotional pictures modulated between-trial pattern similarity in hippocampus and FPl in opposing directions: with higher amygdala activation predicting higher hippocampal similarity across consecutive trials, but lower FPl similarity.

### Within-trial correlation: Higher FPl similarity is associated with poorer temporal memory

Recent work shows that contextual changes (i.e., event boundaries) disrupt neural pattern similarity in temporal-memory sensitive regions even within a trial, reducing within-trial (TR-by-TR) neural pattern correlation (also called temporal autocorrelation), suggesting a role for moment-to-moment pattern similarity dynamics in the temporal organization of memory (Brunec et al., 2018; Kjelstrup et al., 2008; Paz et al., 2010; Raut et al., 2020; Sinclair et al., 2021). Thus, we next tested whether hippocampal and FPl within-trial pattern correlation following the presentation of emotional images were associated with temporal memory in our study. We focused on the third and fourth TRs after image onset, which captured the potential ‘persistence’ of image-related neural activity patterns immediately after image offset (**Figure 2B**; see *Methods* for details).

We found that trials with higher temporal memory performance were associated with lower FPl within-trial pattern correlation between the third and fourth TRs compared to low-performance trials (*p* = 0.037, FDR corrected, **Figure 4A**). Importantly, other TR pairs (during image presentation) did not show this effect (*ps* > 0.468, **Figure 4A**)—suggesting that this was primarily due to the persistence of representational patterns over time. The positive association between within-trial FPl pattern similarity and temporal memory errors was also found when examining temporal memory errors continuously (B = 0.057 (SE = 0.028), F = 4.03, *p* = 0.045, **Figure 4B**). Hippocampal within-trial pattern correlation was not reliably associated with temporal memory errors (B = -0.028 (SE = 0.048), F = 0.33, *p* = 0.563, **Supplementary Figure 2A-B**). Of note, control analyses run using the residual time series did not reveal significant effects (*ps* > 0.279). Collectively, these results suggest that lower within-trial pattern similarity in FPl supports better temporal memory for emotional events.

**Figure 4.**
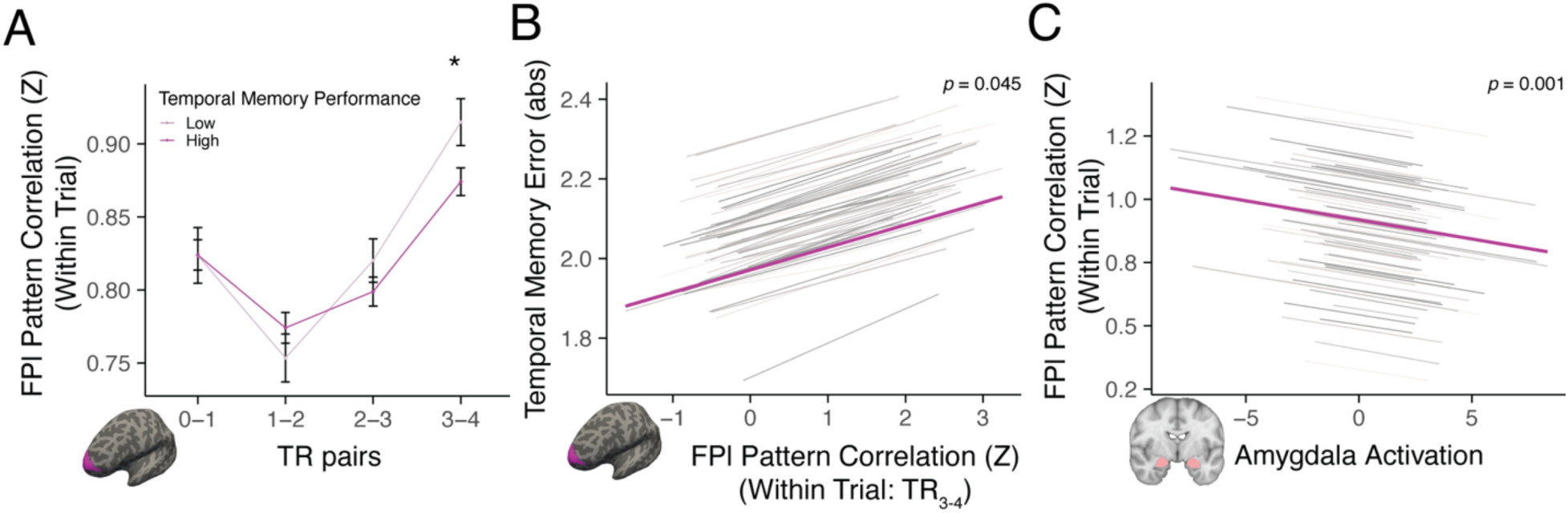
Within-trial pattern correlations. **A-B.** Higher temporal memory performance was associated with lower within-trial FPl pattern similarity (Z_(r:TR 3-4)_) (A), an effect also observed when examining temporal memory errors continuously (B). Error bars: within-subjects standard error for each condition. **C.** Higher amygdala activation was associated with lower within-trial pattern similarity in FPl.

Next, we tested whether amygdala activation modulated within-trial pattern similarity in the FPl. We found that greater amygdalar activation was associated with lower within-trial pattern similarity in the FPl (B = -0.015 (SE = 0.004), F = 11.76, *p* < 0.001, **Figure 4C**).

### Distinct hippocampal and FPl pathways sculpt emotional temporal memory

To recapitulate the above-reported results, negative emotional valence was associated with better temporal memory, and the similarity of neural activity patterns in the hippocampus (between trials) and FPl (within trial) were associated with temporal memory and modulated by amygdalar activation during emotional-picture encoding. Thus, we next examined whether FPl and hippocampal similarity metrics significantly mediated the amygdala-driven impact of emotional valence on temporal memory. To that end, we conducted mediation analyses using Bayesian Regression Models (see *Methods*) whereby we entered between-trial hippocampal and within-trial FPl pattern correlation as mediator of the impact of image valence and amygdala activation on temporal memory—first, as separate models, and then simultaneously. First, we found a significant negative indirect effect of emotional valence via the amygdala-hippocampal pathway on temporal memory (ß = -0.00002, 95% CI = [-0.00005, -0.0000009], *pd* = 98.26%, **Supplementary Figure 3A**), suggesting lower image valence produced higher amygdala activation, which in turn increased hippocampal between-trial pattern correlation, impairing temporal memory. We also found some evidence for a positive indirect effect via the amygdala-FPl pathway (ß = 0.000009, 95% CI = [-0.0000002, 0.00003], *pd* = 97.25%, **Supplementary Figure 3B**), whereby more negative images elicited higher amygdala activation, which in turn decreased FPl within-trial pattern correlation to support better temporal memory. Of note, all correlations between node pairs in the two pathways were significant (*pd* ≥ 97.9%, **Supplementary Figure 3, Supplementary Table 1**), consistent with our above mixed-effects model results.

Given that these mediational models revealed indirect effects of emotion on temporal memory via two distinct amygdala pathways (amygdala-hippocampal and amygdala-FPl), we next conducted an exploratory parallel mediation analysis to test whether both of these pathways still mediated the effect of emotion on temporal memory when considered simultaneously. We found that the level of evidence for both indirect pathways was consistent with the effects reported above (hippocampal between-trial pattern correlation: ß = -0.00002, 95% CI = [-0.00005, -0.0000009], *pd* = 98.38%; FPl within-trial pattern correlation: ß = 0.000008, 95% CI = [-0.0000004, 0.00003], *pd* = 96.54%; **Figure 5**), suggesting that each pathway makes a distinct and independent contribution in mediating how emotional valence sculpts temporal memory. Collectively, these results indicate that emotional valence driven amygdala activation modulates pattern similarity in the hippocampus and FPl at multiple timescales, with the potential to both blur and sharpen temporal memory.

**Figure 5.**
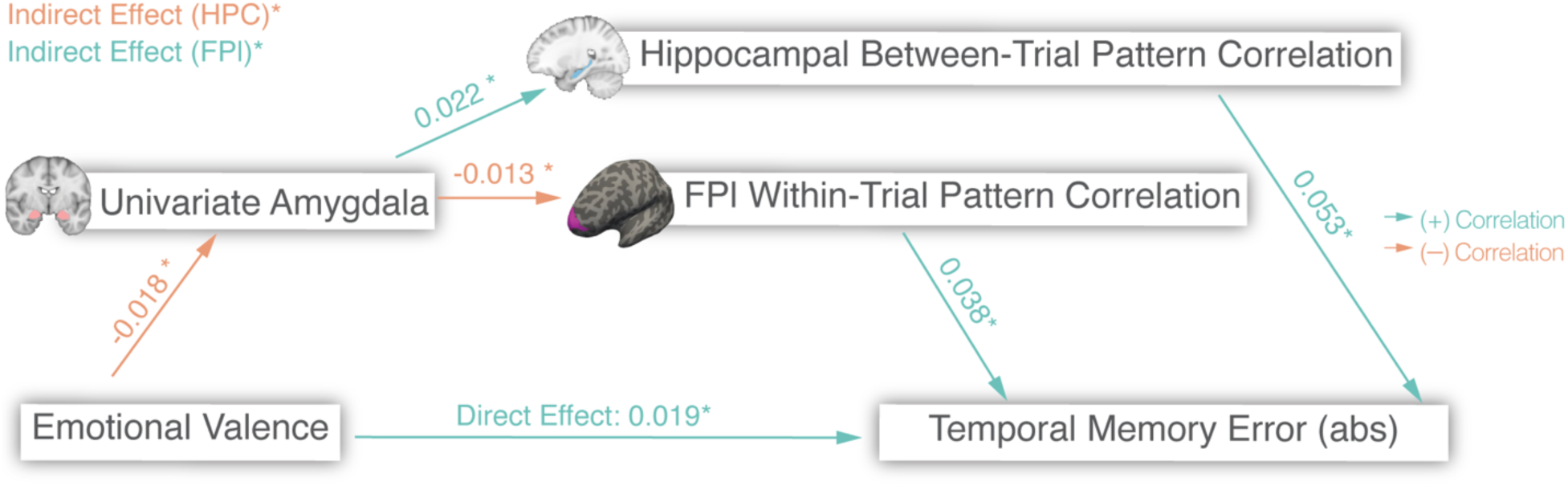
Parallel mediation results indicate that distinct amygdala pathways mediate the influence of emotion on temporal memory. Emotional-valence driven changes in amygdala activation modulated between-trial hippocampal pattern similarity and within-trial FPl pattern similarity, both of which independently mediated the effect of emotional valence on temporal memory. β values between each node in the serial pathway are shown on the arrows. * *pd* value > 95%. Red: negative correlation. Green: positive correlation.

### Perirhinal activation is associated with recency biases in temporal memory estimates

Previous evidence suggests that perirhinal activation during stimulus encoding is associated with recency biases in temporal order judgments (Jenkins & Ranganath, 2016)—an effect that influences temporal memory accuracy as a function of when items were encoded within the session (i.e., correlating with temporal memory accuracy for recently-presented items; (Eichenbaum et al., 2007; Hintzman, 2010; Ranganath & Ritchey, 2012). Therefore, we next examined the association between perirhinal activation at encoding and temporal memory judgments for emotional pictures as a function of the time at which each image was encoded during the session (“session time”).

We found a significant interaction between session time and perirhinal activation on temporal memory accuracy (F = 10.59, *p* < 0.001, **Figure 6A**), such that stronger perirhinal activation at encoding was associated with lower temporal errors only for recently-presented items (i.e., in the “Late” part of the session) (Late: *p* < 0.001, Mid: *p* = 0.175, Early: *p* = 0.450). We tested whether this association was due to perirhinal-related shifts in recency estimates by additionally examining *directional* temporal memory errors, which indexed whether temporal estimates for each image were biased toward the end or towards the beginning of the experimental session. Critically, we found a significant interaction between session time and perirhinal activation when predicting directional temporal memory judgments (F = 11.64, *p* < 0.001, **Figure 6B**), such that higher perirhinal activation was associated with a temporal memory bias toward the end of the session (an effect driven primarily by stimuli shown in the middle and end of the session) (Late: *p* < 0.001, Mid: *p* < 0.001, Early: *p* = 0.434). Collectively, these results indicate that perirhinal activation during emotional-stimulus encoding is associated with recency biases in temporal memory estimates, an effect that differentially drives temporal memory accuracy as a function of encoding recency.

**Figure 6.**
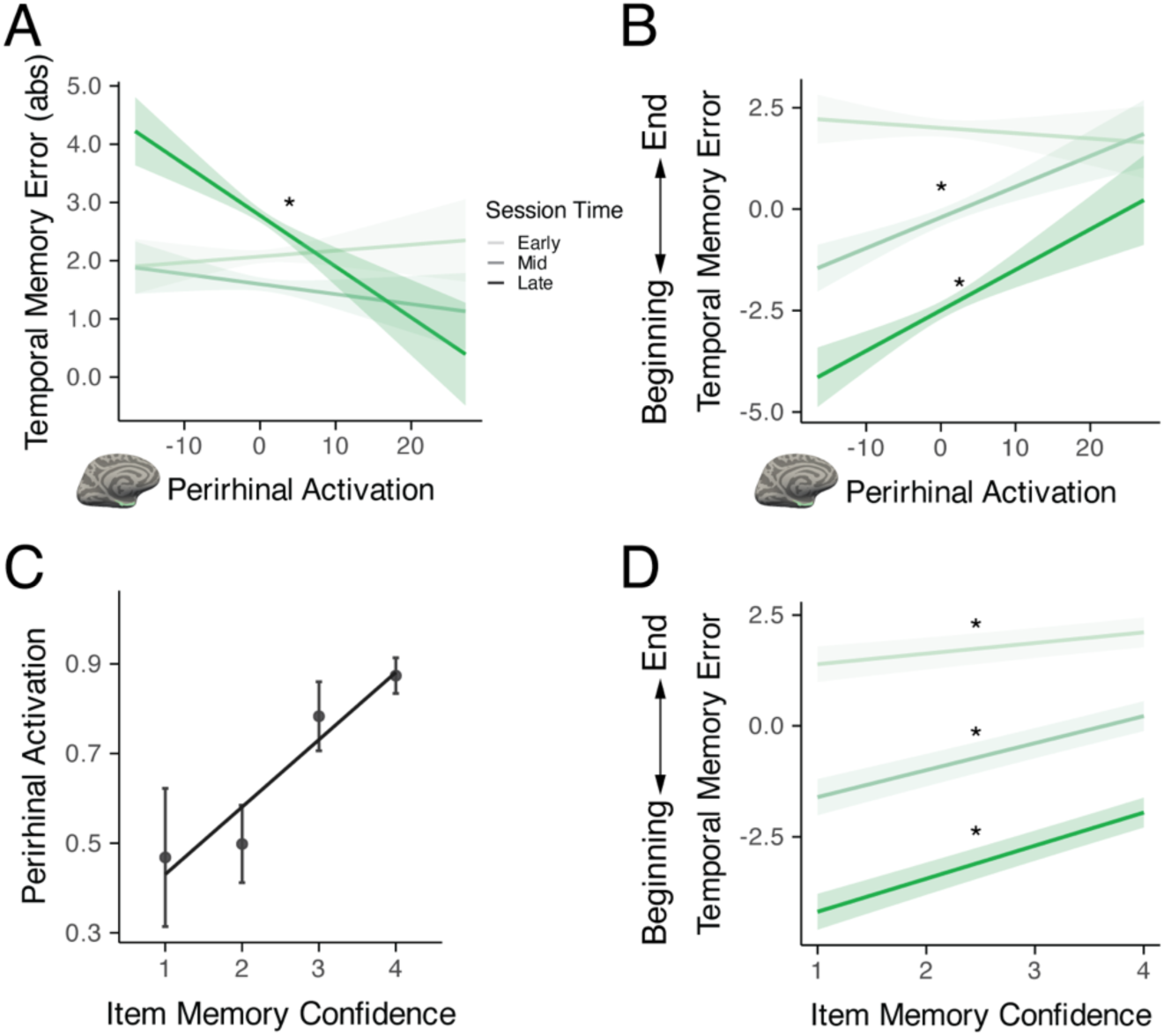
Perirhinal activation predicts recency biases in temporal memory estimates. **A.** Image presentation time (“session time”) and perirhinal activation significantly interact when predicting absolute temporal memory errors, whereby higher perirhinal activation at encoding predicted lower temporal memory errors for recently-presented items. **B.** Analyses of directional errors revealed that perirhinal activation at image encoding was associated with temporal memory estimates biased toward the end of the experiment (an effect that is pronounced for trials occurring in the middle and late portions of the experimental session). **C.** Higher perirhinal activation during encoding was associated with higher confidence in subsequent item memory judgments. **D.** Stimulus presentation time (“session time”) and confidence in item memory judgments interacted when predicting directional temporal memory errors, such that higher item memory confidence predicted a greater bias toward the end of the session for recently-presented items.

Theoretical work has proposed a strength-based model for how perirhinal-dependent representations may drive temporal memory judgments (DuBrow & Davachi, 2017), whereby perirhinal contributions to item memory strength may mediate the effects of perirhinal function on temporal memory. To investigate this hypothesis in our data, we tested the relationship between perirhinal activation and item memory strength as indexed by item memory confidence (Yonelinas, 2023). First, emotional stimuli reported as “remembered” were associated with lower temporal memory error than those reported as “forgotten” (B = -0.191 (SE = 0.018), F = 109.15, *p* < 0.001, **Supplementary Figure 4A**). Critically, images endorsed as “remembered” with higher confidence were associated with larger perirhinal activation (**Figure 6C)** (Remembered: B = 0.147 (SE = 0.027), F = 29.80, *p* < 0.001 *vs*. Forgotten: B = -0.028 (SE = 0.038), F = 0.53, *p* = 0.465). Further, item memory confidence and session time interacted when predicting directional temporal memory biases (F = 37.01, *p* < 0.001, **Figure 6D**), whereby higher item memory confidence was associated with a larger temporal bias toward the end of the session (an effect particularly pronounced for later-occurring events; late *vs*. early: *p* < 0.001, late *vs*. mid *p* = 0.068). Collectively these findings align with a strength-based representational model for temporal memory judgments (DuBrow & Davachi, 2017).

Finally, given established interactions between the amygdala and perirhinal cortex in emotion-modulated memory (Bisby & Burgess, 2017; Yonelinas & Ritchey, 2015), we examined the association between amygdala and perirhinal engagement to emotional images and tested whether emotional valence modulated the association between item memory and perirhinal engagement. We found that emotional images that were more negative (B = -0.099 (SE = 0.01), χ^2^(1) = 96.35, *p* < 0.001, **Supplementary Figure 4B**) and higher in arousal (B = 0.119 (SE = 0.024), χ^2^(1) = 24.43, *p* < 0.001, **Supplementary Figure 4C**) were more likely to be reported as remembered. Images that were more negative also evoked higher perirhinal activation (B = -0.031 (SE = 0.011), F = 7.49, *p* = 0.010, **Supplementary Figure 4D**), which covaried with trial-wise amygdala responses (B = 0.548 (SE = 0.015), F = 1370.93, *p* < 0.001), consistent with prior work (Ritchey et al., 2008, 2019) (for additional analyses, including Bayesian mediation models, see Supplementary Results & **Supplementary Figure 4E**). In summary, our results suggest that negative emotional valence may strengthen perirhinal contributions to memory, shaping both item memory strength and the tendency to judge events as having occurred more recently. Nonetheless, note that trial-wise perirhinal engagement continued to predict recency-biased temporal memory judgments even after controlling for item memory (for this and other analyses controlling for item memory, see *Supplementary Results: Control analyses: item memory*).

## Discussion

Extant work examining the neural bases of temporal memory, conducted primarily in the context of neutral-event processing, shows that changes in the similarity of neural activity patterns in the hippocampus and FPl often reflect the objective passage of time (Buonomano et al., 2023; Hsieh et al., 2014; Nielson et al., 2015) and track subjective temporal memory estimates (Ezzyat & Davachi, 2014; Gauthier et al., 2020; Jenkins & Ranganath, 2010, 2016; Tsao et al., 2022). We found that amygdala engagement in response to emotional pictures modulates distinct neural correlates of temporal memory via parallel prefrontal and medial temporal lobe pathways. Specifically, higher pattern similarity in the hippocampus between successive emotional pictures was associated with poorer accuracy in estimating *when* emotional pictures occurred during the experiment. Greater within-trial similarity between FPl activity patterns during emotional-picture presentation and after picture offset was also associated with poorer temporal memory. Critically, stronger amygdala engagement during emotional-picture encoding was associated with opposing effects in hippocampal *vs*. FPl pattern similarity, with larger amygdala responses predicting higher similarity in hippocampal patterns over time, but lower similarity in FPl patterns—consistent with known neuroanatomical projections. In addition, greater emotional-image induced perirhinal activation at encoding was associated with a recency bias in temporal memory estimates. Collectively, this work reveals distinct amygdala-FPl and amygdala-MTL pathways that sculpt temporal coding of emotional events, with the potential to sharpen and blur temporal memory.

### Between-trial pattern similarity in the hippocampus predicts temporal memory fidelity for emotional events

Multivariate pattern changes in the hippocampus and FPl have been shown to correlate with the passage of time and support temporal context representations (reviewed in (Buonomano et al., 2023; Clewett et al., 2019; Eichenbaum, 2017b; Kwok et al., 2025; Sugar & Moser, 2019; Wang et al., 2022)). For instance, greater hippocampal pattern dissimilarity across distinct events has been linked with longer objective (Hsieh et al., 2014; Mankin et al., 2015; Nielson et al., 2015; Wang et al., 2025) and remembered (Ezzyat & Davachi, 2014) temporal distance for events. Moreover, the similarity of hippocampal patterns over time has also been proposed to relate to the fidelity of temporal coding in memory (Eichenbaum, 2017b; Wang et al., 2022). For instance, greater similarity in the rodent hippocampal place-cell firing patterns between distinct events is related to lower temporal order memory (Manns et al., 2007), consistent with an effect previously observed in human hippocampal BOLD activation patterns (Jenkins & Ranganath, 2016) (but see also (DuBrow & Davachi, 2014)). Higher similarity in FPl patterns evoked by adjacent events has likewise been linked to poorer temporal memory accuracy in humans (Jenkins & Ranganath, 2010). Here, we found that higher between-trial pattern similarity in the hippocampus was associated with lower temporal memory accuracy for when emotional events occurred. Although we observed a trend in the same direction, the relationship between between-trial pattern similarity in the FPl and temporal memory accuracy did not reach statistical significance. Collectively, these findings suggest that more differentiated stimulus-evoked neural activity patterns in the hippocampus support higher-fidelity temporal memory for emotional events.

### Within-trial pattern similarity in the FPl is associated with temporal memory fidelity for emotional events

Recent studies suggest that neural pattern changes at short time scales (e.g., within a trial) are sensitive to changes in context (Rait et al., 2025; Sinclair et al., 2021). For example, higher hippocampal drift rates have been linked to shifts in the context in which events unfold (Rait et al., 2025). Moreover, pattern changes in the human hippocampus (Sinclair et al., 2021) and in the rat entorhinal cortex (Kanter et al., 2025) have been shown to accelerate at event boundaries. These studies suggest that contextual changes can lead to neural pattern changes on the scale of seconds, which may support fine-grained temporal coding (Eichenbaum, 2017b; Wang et al., 2022). Here, we found that lower within-trial pattern correlation in FPl was associated with higher-fidelity temporal estimates for when emotional events occurred. This effect was directionally consistent with the above-reported association of temporal memory and between-trial pattern correlation. Of note, our within-trial pattern similarity metric that correlated with temporal memory indexed the persistence of image-evoked neural patterns following stimulus offset (i.e., during the inter-trial interval). The finding that FPl within-trial similarity is associated with finer grained temporal coding extends previous work on the role of FPl in temporal memory (Jenkins & Ranganath, 2010) and aligns with a broader literature documenting a role for FPl in temporally extended control (Desrochers et al., 2015; Nee & D’Esposito, 2017).

### Amygdalar-mediated hippocampal and FPl pathways differentially predict temporal memory

We found that larger amygdala responses to emotional events were associated with higher between-trial hippocampal pattern correlation, consistent with a neuroanatomically inspired theoretical framework on emotion-modulated temporal memory (Wang et al., 2022). The amygdala sends direct and powerful inputs to the upper layers of the hippocampal CA fields, which receive entorhinal inputs thought to support the integrity of hippocampal temporal coding (Howard et al., 2014; Robinson et al., 2017; Saunders & Rosene, 1988; Zeidman & Maguire, 2016). This connectivity pattern can therefore give rise to a competition between amygdala and entorhinal projections within the hippocampus (Wang & Barbas, 2018)—and a push-and-pull between emotion and temporal coding in this region (Wang et al., 2022). Moreover, amygdala inputs can excite parvalbumin inhibitory interneurons, thereby potentially suppressing pyramidal neurons in the hippocampus that receive entorhinal inputs (Wang & Barbas, 2018). Thus, amygdala-hippocampal terminations have the potential to disrupt contextual representation processes in the hippocampus (John et al., 2024)—and be detrimental to temporal coding in the hippocampus (Wang et al., 2022). Consistently, our findings show that amygdala responses are associated with less discriminable hippocampal patterns across discrete time points spanning successive, distinct events.

In contrast, we found that stronger emotional-picture-induced amygdala activation was associated with lower pattern similarity in the FPl (within and between-trials), a finding that aligns with existing anatomical evidence (Ghashghaei et al., 2007; Joyce & Barbas, 2018). Specifically, amygdalar inputs are thought to reach FPl in part via vmPFC, including BA25 and BA32 (Joyce & Barbas, 2018; Medalla & Barbas, 2010). These vmPFC inputs often terminate in the upper layers of FPl and preferentially target calbindin inhibitory neurons, a connection pattern consistent with a feedback system (Barbas, 2015; Barbas et al., 2018). Such feedback circuits are thought to sharpen representations in downstream areas (Keller et al., 2012; Marques et al., 2018), supporting fine-grained representations. Consistently, human neuroimaging studies show that stronger coupling between BA25 and the FPl during negative emotional processing predicts stronger multivariate decoding of emotional valence in the FPl (Lapate et al., 2022), suggesting that BA25 inputs support discriminable emotional context representations in FPl. In addition, stronger structural connectivity of the amygdalofugal pathway connecting the amygdala and FPl has been found to be associated with stronger emotion-driven task performance (Bramson et al., 2020), further supporting that the amygdala–FPl pathway may enhance emotional behavior that requires contextual representations. Consistently, we found that higher amygdala activation in response to emotional images was associated with lower within-trial pattern correlation in the FPl, which was in turn linked to better temporal memory. In summary, these distinct anatomical connectivity profiles may underlie differential amygdala influences on hippocampal and FPl pattern similarity, which in turn can differentially shape temporal memory by blurring or sharpening temporal representations, respectively.

### Two distinct pathways supporting opposing effects of emotion on temporal memory

Consistently, Bayesian mediation analyses indicated that emotional valence influenced temporal memory through both the amygdala-hippocampal and amygdala-FPl pathways with opposing effects. Specifically, more negatively valenced images produced higher amygdala activation, which increased between-trial hippocampal pattern similarity to impair temporal memory, but decreased within-trial FPl pattern similarity to support better temporal memory. These distinct pathways culminated in potentially opposing net impacts of emotional responses on temporal memory and recapitulate the mixed results from a surge of recent behavioral studies, which underscore that emotional events can sometimes impair (Clewett & McClay, 2024; Colson et al., 2025; Cui et al., 2023; Hennings et al., 2021; Laing & Dunsmoor, 2025; Laing et al., 2026; Maddock & Frein, 2009; McClay et al., 2023; Wang & Lapate, 2025; Zlomuzica et al., 2016) and sometimes enhance (Bogdan et al., 2023; Clewett & McClay, 2024; Cliver et al., 2024; D’Argembeau & Van der Linden, 2005; Dev et al., 2022; Lake et al., 2016; McClay et al., 2023; Riegel et al., 2023; Rimmele et al., 2012; Schmidt et al., 2011; Yick et al., 2015) temporal memory. In our study, participants showed higher temporal memory accuracy for more negatively valenced images, as well as for images with lower arousal levels. These opposing, and sometimes interacting, contributions of valence and arousal dimensions may partially account for the mixed findings previously observed in the literature of emotion-temporal memory interactions. It is important to note, however, that the temporal memory metric adopted in this study (a timeline task) is thought to capture relatively coarse temporal memory representations (Jenkins & Ranganath, 2010), which are more often enhanced (*vs*. impaired) by negative emotion (Palombo et al., 2021; Petrucci & Palombo, 2021; Rimmele et al., 2011)—compared to other types of temporal memory, such as temporal order memory (Huntjens et al., 2015; McClay et al., 2025). Future work incorporating a broader range of temporal memory measures—including tasks that permit higher-precision estimates—will be required to fully characterize emotional influences on temporal memory and its underlying neural pathways, and to test whether these effects and putative neural mechanisms vary as a function of the precision demands of the temporal memory task.

### Perirhinal activation at encoding predicts recency biases in temporal estimates

While previous studies have implicated perirhinal activation during encoding in enhanced temporal memory for neutral events (Hannesson et al., 2004; Montchal et al., 2019), other work suggests that this association is explained in part by recency biases in temporal memory judgments associated with greater perirhinal engagement at stimulus encoding (DuBrow & Davachi, 2017; Jenkins & Ranganath, 2016). Here, we replicated and extended these effects, whereby perirhinal engagement during emotional-picture processing was associated with temporal memory accuracy only for recently-encoded images. This finding is consistent with a perirhinal role in familiarity-based recognition, which often shows a recency effect (Howard Eichenbaum, 2017a; Hintzman, 2004, 2010).

Moreover, we found that higher perirhinal activation biased temporal judgments toward the end of the experiment, especially for events encoded later in the session, in alignment with prior work (Jenkins & Ranganath, 2016). A memory strength-based mechanism has been proposed to account for these effects (Yonelinas, 2023), whereby perirhinal-driven increases of item memory strength result in a bias toward judging items as having occurred more recently. Supporting this account, we found that higher item memory confidence—a proxy for item memory strength (Yonelinas, 2023)—was associated with temporal memory judgments biased toward the end of the experiment. Replicating and extending a large body of work (Aly et al., 2013; Bisby & Burgess, 2017; Bisby et al., 2016; Kensinger & Corkin, 2003; LaBar & Cabeza, 2006; Mather & Sutherland, 2009; Ritchey et al., 2019), we found that images with lower emotional valence (i.e., more negative) produced higher perirhinal activation, which in turn correlated with increased memory confidence in remembered trials. A mediation model further suggested that a strength-based mechanism underlay recency judgments for items, whereby negative emotional stimuli enhanced perirhinal engagement, strengthening item representations which, in turn, biased temporal judgments toward later session times. Collectively, these findings point to a perirhinal-mediated, strength-based mechanism through which negative emotion biases temporal memory, particularly for recent events.

## Limitations

The following limitations of the present work warrant additional investigation. First, our sample included a higher proportion of female than male participants; future work will therefore be needed to fully assess the generalizability of these effects across genders. Second, negative images in the present study were, on average, higher on arousal compared to positive stimuli. Although we controlled for arousal in models examining the effects of emotional valence on temporal memory and neural activity, future work should match and/or manipulate arousal levels across positive and negative stimulus sets to fully disentangle their distinct, and potentially interactive, contributions to temporal memory (McClay et al., 2023). Third, while we controlled for semantic similarity between images using a deep neural network approach (Riberto et al., 2022, 2019), future work systematically varying stimulus category across positive and negative valence stimuli would be ideal to isolate the contribution of semantic content from that of emotional valence category to temporal memory.

In conclusion, our results show that emotional valence modulates temporal memory via distinct amygdala-hippocampal and amygdala-frontopolar pathways, offering new insights into the mechanisms through which emotion shapes memory for *when* events occurred.

## Acknowledgements

This research was funded by NIH grant MH134000 (R.C.L.) and by Aligning Science Across Parkinson’s ASAP-020-519 through the Michael J. Fox Foundation for Parkinson’s Research (S.T.G. and R.C.L.). The authors thank Taylor Li and Kiana Sabugo for their help with data collection.

## Data and code availability

The behavioral and neural data are publicly available on OpenNeuro (https://openneuro.org/datasets/ds005265/versions/1.0.0). Analysis code, masks, and images used in this study are available in the following GitHub repository: https://github.com/LEAPNeuroLab/ASAP/tree/main/Approach%20Avoidance. The data, code, protocols, and key lab materials used and generated in this study are listed in a Key Resources Table alongside their persistent identifiers at https://doi.org/10.5281/zenodo.24487332.

## Supplementary Results

### Confidence rating analyses

The distribution of confidence ratings for item and temporal memory judgments is shown in **Supplementary Figure 5A**. Participants reported moderate confidence in their temporal memory estimates (*M* = 2.27, *SD* = 0.94), with most responses in the “2: Slightly Confident” (33.56%) and “3: Moderately Confident” (31.80%) categories. In contrast, participants reported high confidence in item memory judgments (mean = 3.34, SD = 0.92), with most responses falling in the “4: Very Confident” (60.92%) category (**Supplementary Figure 5B**).

### Perirhinal activation and recency biases: further testing a strength-based model

The results reported in the main manuscript (see: “Perirhinal activation is associated with recency biases in temporal memory estimates”) suggest that increasingly negative events elicited perirhinal engagement, potentially strengthening item memory and producing recency biases in temporal memory judgments. To formally test this model, we conducted an exploratory modulated-mediation analysis using Bayesian Regression Models (see *Methods* for details). We found a significant negative indirect effect for trials occurring late in the session (ß = -0.0002, 95% CI = [-0.0004, -0.00003], *pd* = 98.88%, **Supplementary Figure 4E**), whereby more negative images elicited greater perirhinal activation, leading to increased item memory strength, which in turn produced temporal estimates biased toward the end of the experiment. This effect was significantly stronger late *vs*. early in the session (Late: ß = -0.00007, 95% CI = [-0.0001, -0.000009], *pd* = 98.88%), and *vs*. mid-session (ß = -0.0002, 95% CI = [-0.0003, -0.00002], *pd* = 98.88%). In summary, these results indicate that negative emotional valence strengthens perirhinal contributions to memory, shaping both item memory strength and the tendency to judge events as having occurred more recently—particularly for events processed later in the session.

### Control analyses: item memory

To ensure that the reported relationships between univariate and multivariate neural measures and temporal memory were not driven by item memory alone, we included item memory as a covariate in our mixed-effects models. We found that hippocampal between-trial pattern correlation (trial lag 1) remained positively correlated with temporal memory errors while controlling for item memory (B = 0.082 (SE = 0.036), F = 5.11, *p* = 0.024). After controlling for item memory, FPl within-trial pattern correlation showed a trend-level association with temporal memory errors (B = 0.055 (SE = 0.028), F = 3.57, *p* = 0.053). Finally, perirhinal activation continued to significantly interact with session time in predicting both temporal memory errors (B = 0.004 (SE = 0.011), F = 9.75, *p* = 0.0005) and directional temporal memory errors after controlling for item memory (B = -0.061 (SE = 0.013), F = 10.25, *p* < 0.001). After controlling for item memory, amygdala activation remained a significant predictor of between-trial pattern correlation in the hippocampus (B = 0.021 (SE = 0.004), F = 32.70, *p* < 0.001) and FPl (B = -0.014 (SE = 0.005), F = 8.85, *p* = 0.003), within-trial FPl pattern correlation (B = -0.014 (SE = 0.004), F = 10.36, *p* = 0.001), and perirhinal activation (B = 0.542 (SE = 0.015), F = 1335.18, *p* < 0.001).

## Supplementary Figures

**Supplementary Figure 1.**
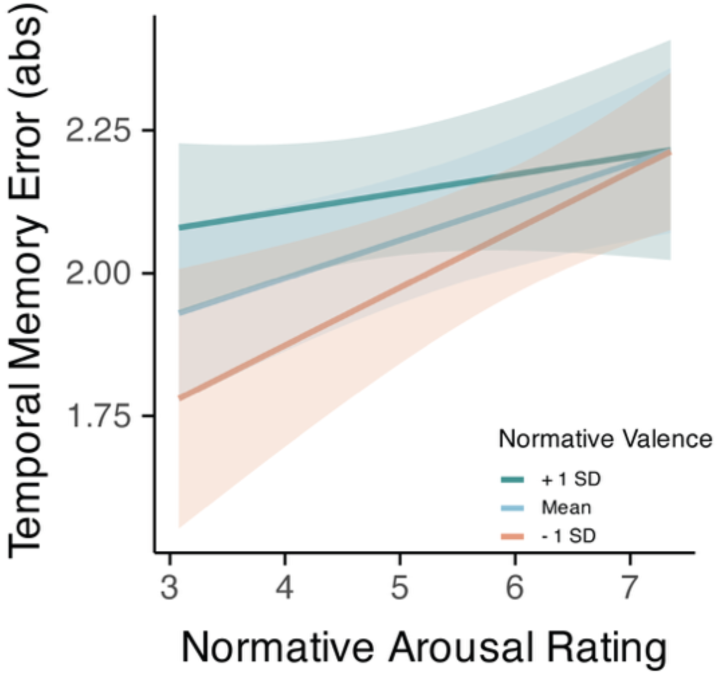
Mixed-effects model showed a trend-level interaction between image valence and arousal when predicting absolute temporal memory errors, whereby the correlation between arousal and temporal memory errors is stronger for pictures with more negative emotional valence. Shaded ribbon in B: 95% confidence interval (CI).

**Supplementary Figure 2.**
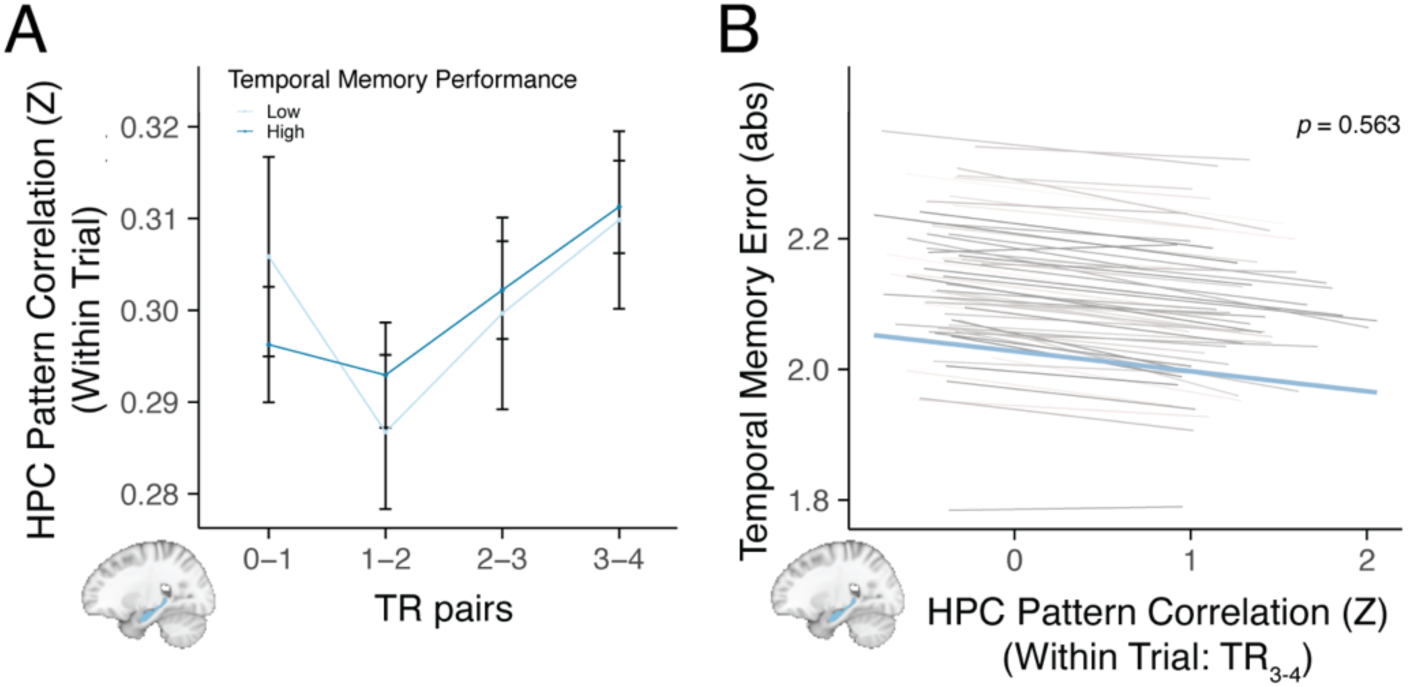
Within-trial pattern correlation in the hippocampus. **A.** Within-trial pattern correlation in the hippocampus did not vary as a function of temporal memory performance. Error bars: within-subjects standard error for each condition. **B.** Likewise, when examined continuously, within-trial pattern correlation (TR3 and TR4 pair) in the hippocampus did not correlate with absolute temporal memory error.

**Supplementary Figure 3.**
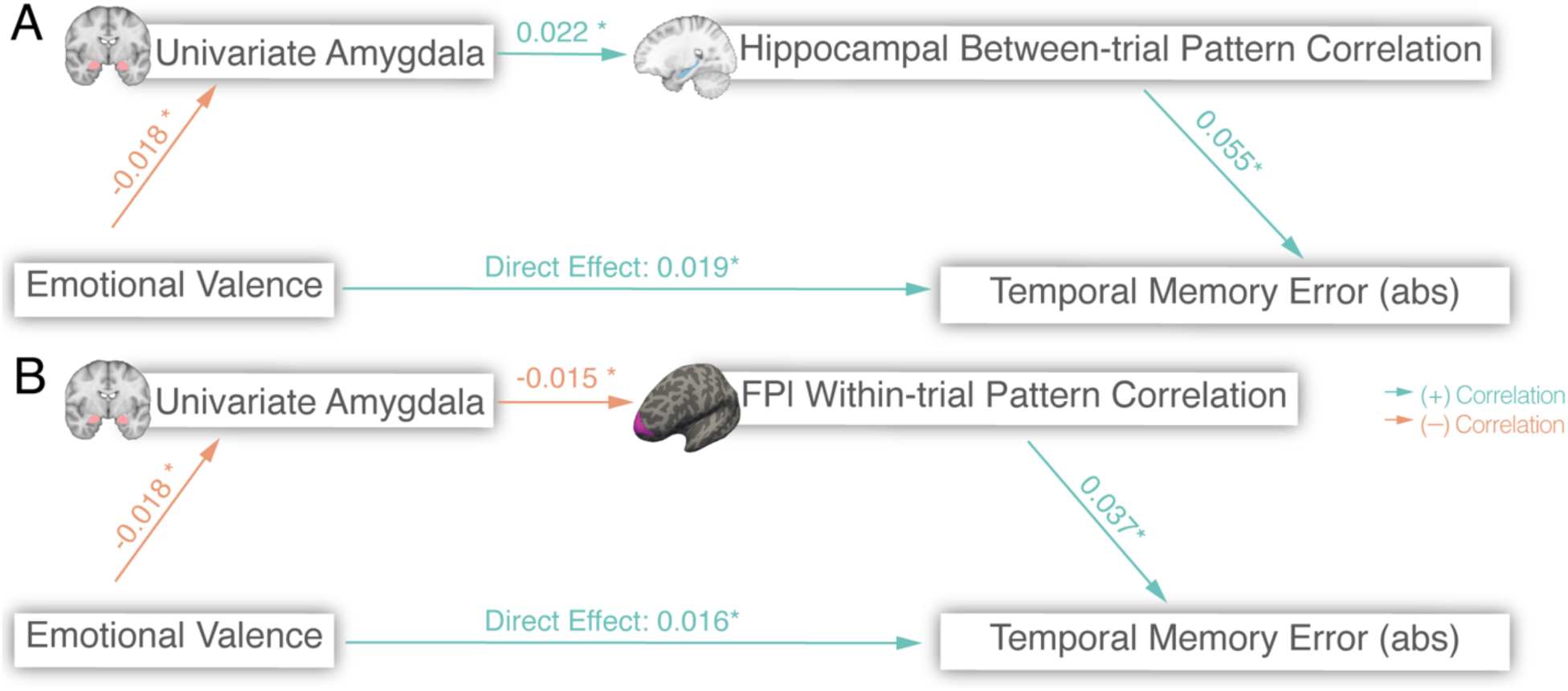
Separate mediational models are shown for amygdala-hippocampal (A) and amygdala-FPl pathways (B) that each mediate the effect of emotional valence on temporal memory. β values between each node in the serial pathway were shown on the arrows. * *pd* value > 95%

**Supplementary Figure 4.**
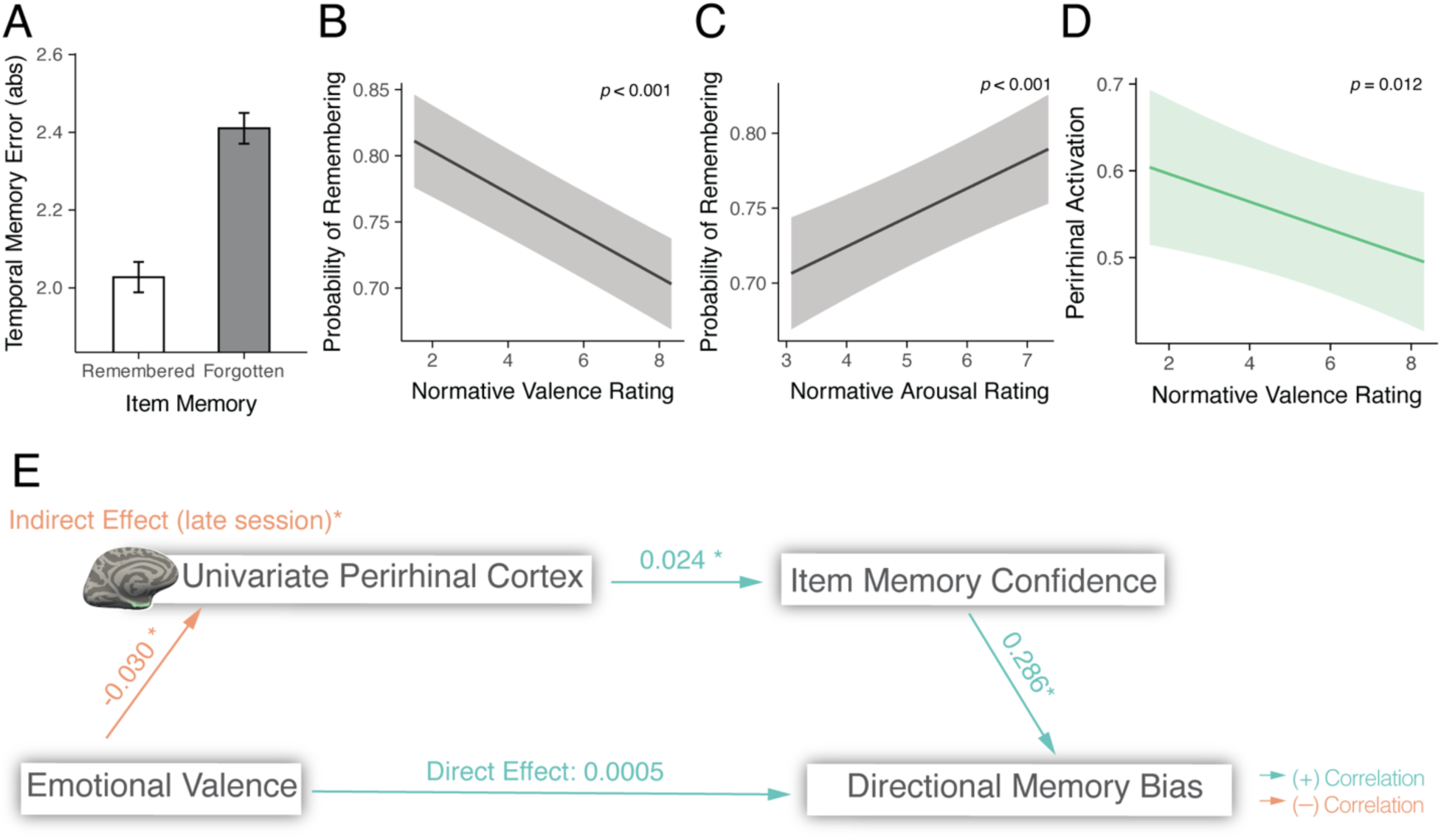
Perirhinal activation predicts recency biases in temporal estimates. **A.** Images endorsed as remembered were associated with lower temporal memory errors than images endorsed as forgotten. **B-C.** Mixed-effects model indicated that lower normative valence ratings (B) and higher arousal ratings (C) were associated with a higher probability of endorsing an image as remembered. **D.** Mixed-effects model indicated that lower valence (i.e., more negative) images were associated with higher perirhinal activation. **E.** Modulated serial mediation results showed that perirhinal activation influenced item memory confidence, which mediated the effect of valence on directional temporal memory estimates for images shown later in the session. β values between each node in the serial pathway were shown on the arrows. * *pd* value > 95%. Shaded ribbon in B-D: 95% confidence interval (CI).

**Supplementary Figure 5.**
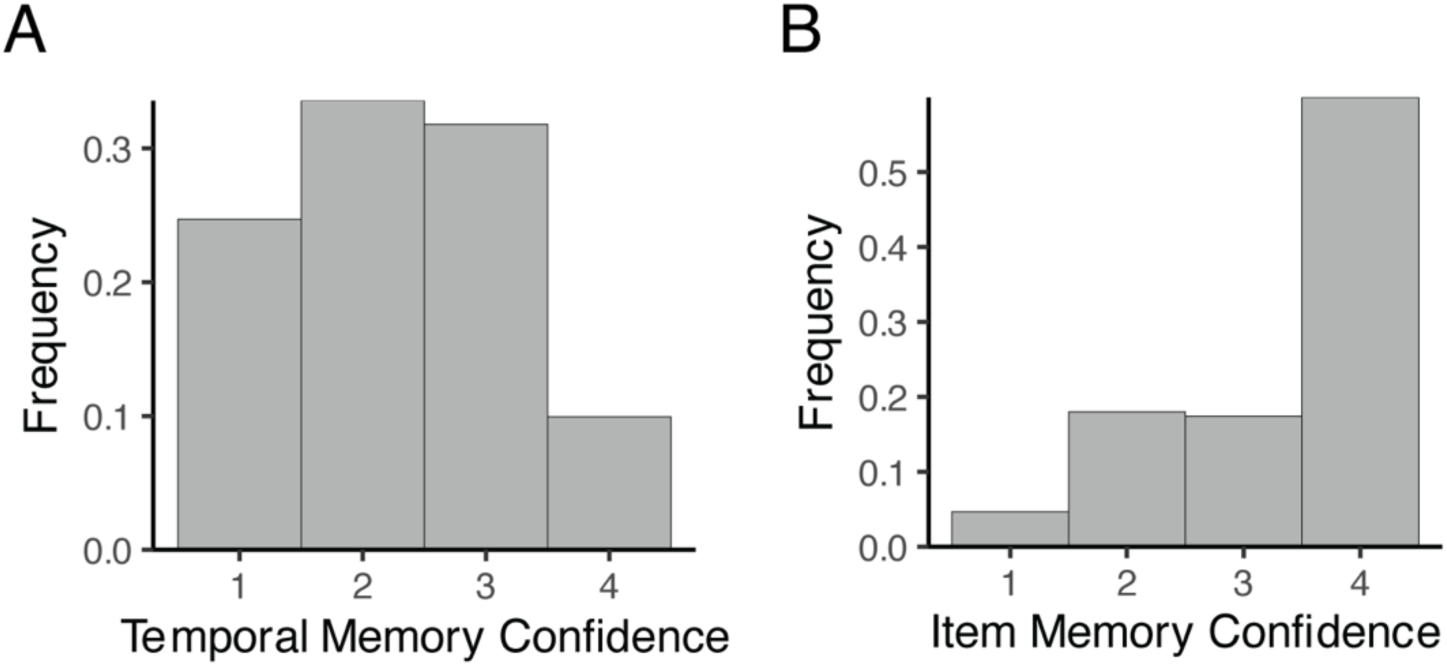
Histograms showing the frequency of confidence ratings given for temporal memory (A) and item memory (B) judgments.

## Supplementary Tables

**Supplementary Table 1.** Statistics of the mediational models.

| **Mediation Model** | **Node Path** | **β** | **CI (95%)** | ***pd*** |
|---|---|---|---|---|
| Amygdalar-Hippocampal Pathway | Valence ->Amygdala | -0.018 | [-0.031, -0.004] | 0.993 |
|  | Amygdala->Hippocampus | 0.022 | [0.014, 0.030] | 1.000 |
|  | Hippocampus->Temporal Memory Error | 0.055 | [0.008, 0.101] | 0.990 |
| Amygdalar-FPl Pathway | Valence->Amygdala | -0.017 | [-0.031, -0.003] | 0.994 |
|  | Amygdala->FPl | -0.015 | [-0.024, -0.006] | 0.999 |
|  | FPl->Temporal Memory | 0.037 | [0.001, 0.073] | 0.979 |

Terms and abbreviations: amygdala: amygdala activation, FPl: within-trial pattern correlation in lateral frontal pole, hippocampus: between-trial pattern correlation in the hippocampus, *pd*: probability of direction.

